# Hepatocyte differentiation requires anisotropic expansion of bile canaliculi

**DOI:** 10.1101/2024.02.19.581065

**Authors:** Lenka Belicova, Maarten Bebelman, Elzbieta Gralinska, Tobias Jumel, Aparajita Lahree, Andrej Shevchenko, Timofei Zatsepin, Yannis Kalaidzidis, Martin Vingron, Marino Zerial

**Author notes:** Author for correspondence L. Belicova’s present address is Karolinska Institutet, Stockholm, Sweden. These authors contributed equally to this work (Lenka Belicova, Maarten Bebelman).

## Abstract

During liver development, bipotential progenitor cells called hepatoblasts differentiate into hepatocytes or cholangiocytes. Hepatocyte differentiation is uniquely associated with multi-axial polarity, enabling the anisotropic expansion of apical lumina between adjacent cells and formation of a three-dimensional network of bile canaliculi (BC). Cholangiocytes, the cells forming the bile ducts, exhibit the vectorial polarity common to epithelial cells. Whether and how cell polarization feeds back on the gene regulatory pathways governing hepatoblast differentiation is unknown. Here, we used primary hepatoblasts to investigate the contribution of anisotropic apical expansion to hepatocyte differentiation. Silencing of the small GTPase Rab35 caused isotropic lumen expansion and formation of multicellular cysts with the vectorial polarity of cholangiocytes. Gene expression profiling revealed that these cells express reduced levels of hepatocyte markers and upregulate genes associated with cholangiocyte identity. Time-course RNA sequencing demonstrated that loss of lumen anisotropy precedes these transcriptional changes. Independent alterations in apical lumen morphology induced either by modulation of the subapical actomyosin cortex or increased intraluminal pressure caused similar transcriptional changes. These findings suggest that cell polarity and lumen morphogenesis feedback to hepatoblast-to-hepatocyte differentiation.

**Summary statement:** Differentiation of liver progenitors to functional hepatocytes requires anisotropic elongation of their nascent apical surfaces into tubular bile canaliculi.

## Introduction

Cellular differentiation and morphogenesis occur in concert to build functional tissues during development. Whereas differential gene expression determines the building blocks that shape cells and tissues, tissue morphogenesis conversely feeds back on gene regulatory pathways and impacts cell fate decisions via external chemical and mechanical cues (Tatapudy et al., 2017, Motegi et al., 2020, Haftbaradaran Esfahani and Knoll, 2020, Chan et al., 2017). The developing liver provides an excellent example to explore the intertwined mechanisms regulating cell fate. Hepatoblasts are bipotential liver progenitors that differentiate into two functionally and morphologically distinct epithelial cell types: hepatocytes and cholangiocytes (Musch, 2018, Tanimizu and Mitaka, 2017). Hepatocytes are liver parenchymal cells with an unusual cell polarity that is unique among epithelia (Morales-Navarrete et al., 2019). In contrast to simple epithelia which have a single apico-basal axis, hepatocytes have multiple apical (and basal) surfaces. Their apical surfaces are shared between juxtaposed hepatocytes and grow anisotropically i.e., forming elongated tubules instead of spheres. These fine tubules, known as bile canaliculi (BC), form a network where bile is secreted and transported to bile ducts, larger epithelial tubes formed by cholangiocytes that transfer bile to the intestine. Contrary to hepatocytes, cholangiocytes display the common vectorial polarity, whereby their apical surfaces face a common lumen, similar to epithelial tubes in other organs.

We recently described that to form BC, hepatocytes generate transversal apical connections, termed apical bulkheads, that mechanically connect the apical membranes of juxtaposed hepatocytes and enforce the anisotropic expansion of nascent apical lumina into tubular BC (Bebelman et al., 2023, Belicova et al., 2021). Loss of apical bulkheads, through genetic manipulation (see below), results in the isotropic expansion of newly formed apical lumina leading to the formation of multicellular cysts in which the cells adopt a vectorial polarization, similar to cholangiocytes.

During development, the vast majority of hepatoblasts differentiate into hepatocytes through a default differentiation program, while hepatoblasts near the portal veins branch away from the default program to differentiate into cholangiocytes by suppressing the hepatocyte-specific and activating a cholangiocyte-specific transcriptional program (Yang et al., 2017, Prior et al., 2019, Yang et al., 2023). The timing of hepatocyte differentiation (E14.5-E15.5) coincides with hepatoblast polarization and BC formation (Delpierre et al., 2024). How hepatoblast polarization and differentiation are linked and contribute to hepatocyte cell fate determination is an unexplored problem. Recent studies in a variety of cellular and animal model systems have revealed a functional link between the molecular mechanisms underlying cell polarization, mechanical forces at the cell and tissue level, and the genetic program of cell lineage in epithelia (Rose and Gonczy, 2014, Lamba and Zernicka-Goetz, 2023, Ahuja and Cleaver, 2022, Motegi et al., 2020, Hashimoto and Munro, 2018, Hirate et al., 2013).

Whether and how the cellular polarization phenotype feeds back on the gene regulatory pathways that govern hepatoblast differentiation is currently unclear. We have recently modelled hepatocyte differentiation in the embryonic liver by *in vitro* culture of primary mouse hepatoblasts. Using this system, we identified a key regulator, the small GTPase Rab35, that plays a role in actin organization and endocytic recycling (Klinkert and Echard, 2016) and whose expression can be manipulated to switch the polarity phenotype of the differentiating cells (Belicova et al., 2021). Instead of the typical multi-axial polarization of hepatocytes, upon silencing of Rab35, hepatoblasts adopt a vectorial polarization with their apical membranes facing the central lumen, similar to cholangiocytes in bile ducts. We took advantage of this system to examine the consequence of a switch from multi-axial to vectorial polarity for the cell fate determination of hepatoblasts.

## Results and discussion

### Rab35 silencing impairs hepatocyte polarization and differentiation

During the *in vitro* differentiation of primary hepatoblasts towards hepatocytes, nascent apical lumina between adjacent cells expand anisotropically to form elongated BC (Fig. 1A), recapitulating BC formation *in vivo* (Belicova et al., 2021). Silencing of the small GTPase Rab35 results in a loss of apical bulkheads, isotropic apical lumen expansion, and ultimately, the formation of multicellular cysts (Fig. 1A). We first verified that this phenotype is not an epiphenomenon but reflects the requirement of a molecular pathway required for apical bulkheads formation. We therefore tested whether silencing of the Rab35 guanine-nucleotide exchange factor DENN domain containing 1B (Dennd1b), or the Rab35 effector ArfGAP with coiled-coil, ankyrin repeat and PH domains 2 (Acap2), can phenocopy the loss of Rab35. Indeed, their silencing also resulted in isotropic lumen expansion and cyst formation (Fig. 1A, Fig. S1A), indicating the requirement of a Rab35 pathway for the polarity of hepatocytes.

**Figure 1.**
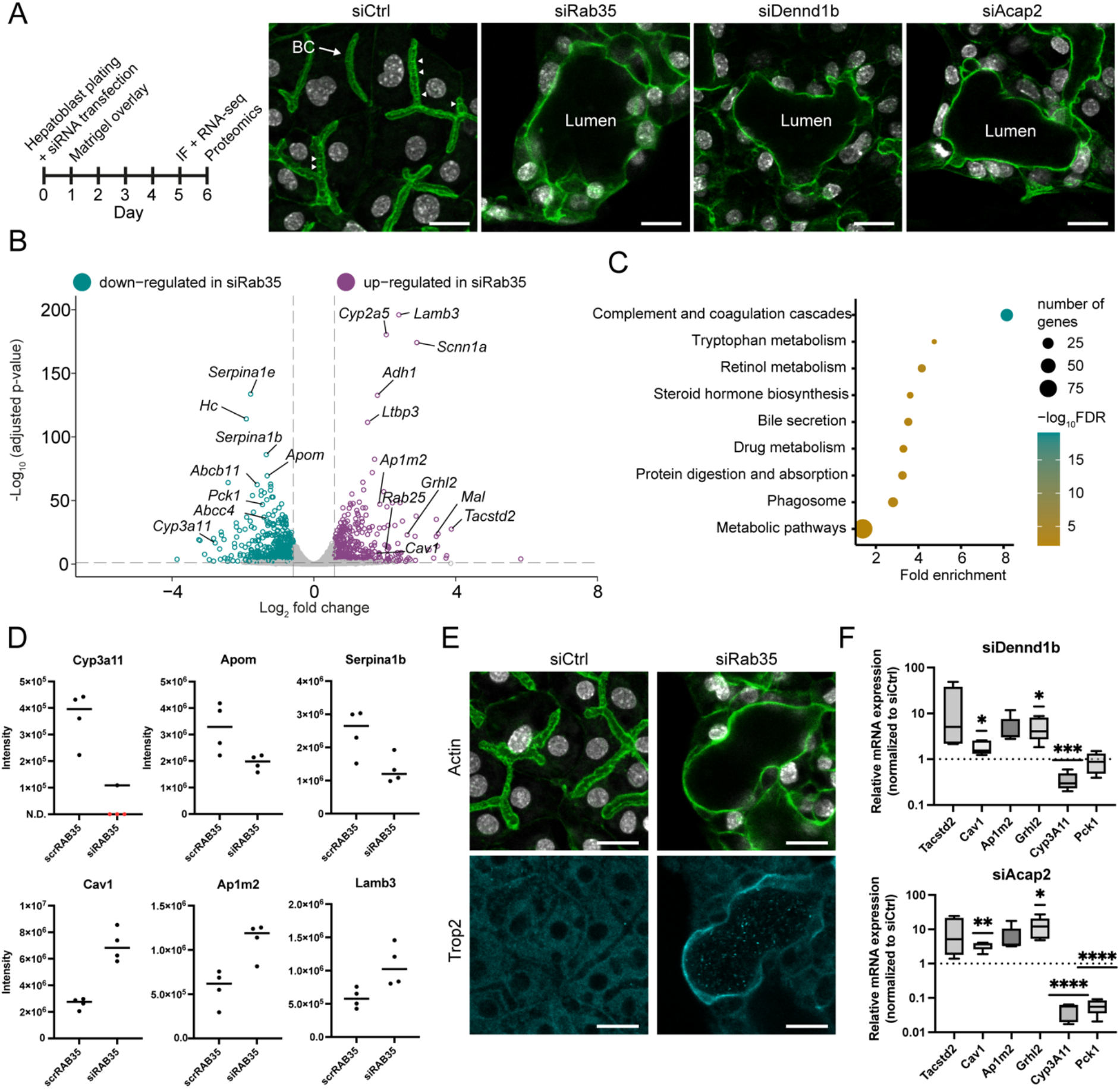
Rab35 silencing impairs hepatocyte polarization and differentiation. A) Left: Design of the hepatoblast culture protocol, siRNA-transfection, and analysis methods. IF: Immunofluorescence microscopy. Right: Hepatoblasts differentiating into hepatocytes in culture transfected with control siRNA (siCtrl) or siRNAs targeting Rab35, Dennd1b and Acap2. F-actin (green), nuclei (grey). Arrowheads highlight several apical bulkheads. B) Volcano-plot showing differentially regulated genes upon siRab35-transfection, compared to siCtrl. C) Selected pathways from KEGG Pathway enrichment analysis based on the downregulated genes (772) in siRab35 transfected cells. D) Mass spectrometry-based MaxLFQ intensities of proteins associated with hepatocyte identity (Cyp3a11, Apom, Serpina1b) and vectorial polarity and cholangiocyte identity (Cav1, Ap1m2, Lamb3). N.D: Not detected. E) SiCtrl- or siRab35-transfected differentiating hepatoblasts. Trop2 (cyan), F-actin (green), nuclei (grey). F) qRT-PCR for genes associated with vectorial polarity and cholangiocyte identity (*Tacstd2, Cav1, Ap1m2, Grhl2*) and hepatocyte identity (*Cyp3a11, Pck1*) following siDennd1b and siAcap2-transfection. Box plots represent 5 independent experiments. *, P < 0.05; **, P < 0.01. Scalebars: 20 μm.

Given the emerging relationship between polarization, mechanical forces and the genetic program of cell lineage in epithelia, the change of cell polarity caused by Rab35 knockdown raises the question as to what extent hepatoblast-to-hepatocyte differentiation is also affected. Previously, we showed that cells in the cysts still expressed the hepatocyte marker HNF4α but were negative for the cholangiocyte marker Sox9 (Belicova et al., 2021), suggesting that they may retain their hepatocyte identity. However, to explore their differentiation state more thoroughly and comprehensively, we used RNA sequencing to profile gene expression of cells treated with control (luciferase) and Rab35-targeting siRNAs at day 5 of the culture, when the morphological phenotype is well pronounced (Fig. 1A). We identified 1261 differentially expressed genes (log2 fold change > |0.58|, adjusted p-value < 0.01) (Fig. 1B). Interestingly, amongst the downregulated genes in siRab35-transfected cells, we found several typical hepatocyte-associated genes, such as *Cyp3a11, Serpina1b/e, Pck1* and *Apom* (MacParland et al., 2018, Liang et al., 2022, Prior et al., 2019). Furthermore, KEGG Pathway enrichment analysis showed that the downregulated genes (772) were enriched in pathways typical for mature hepatocyte function (Fig. 1C, Table S1). In contrast to the downregulation of hepatocyte-specific genes, we observed significant upregulation of several genes typically not expressed in hepatocytes but related to vectorial polarity and cholangiocyte identity, e.g., *Ap1m2, Mal, Cav1, Rab25, Lamb3, Grhl2 and Tacstd2* (Fig. 1B) (Senga et al., 2012, Yamada et al., 2020, Ramnarayanan et al., 2007, Folsch, 2015, Segal et al., 2019, Aizarani et al., 2019, Yang et al., 2017).

To verify that changes in gene expression upon Rab35 knockdown were also reflected at the protein level, we performed proteomic analysis on cell lysates collected at day 6, allowing sufficient time for translation of upregulated mRNAs into proteins. Consistent with the RNA sequencing results, Rab35 knockdown reduced the expression of several proteins associated with hepatocyte fate and function, e.g., Cytochrome P450 3A11 (*Cyp3a11*), Alpha-1-antitrypsin 1-2 (*Serpina1b*), Apolipoprotein M (*Apom*). Conversely, it increased the expression of several proteins associated with cholangiocyte identity and vectorial polarity, e.g. Caveolin-1 (*Cav1*), AP-1 complex subunit mu-2 (*Ap1m2*), and Laminin β3 (*Lamb3*) (Fig. 1D). Furthermore, immunofluorescence microscopy revealed expression of Trop2 (also known as Tumor-associated calcium signal transducer 2, *Tacstd2*) and Caveolin-1 (*Cav1*) specifically in the cells forming cysts (Fig. 1E, Fig. S1B). Similar to the silencing of Rab35, knockdown of Dennd1b or Acap2 decreased *Cyp3a11* and *Pck1* mRNA levels, while increasing *Ap1m2, Cav1, Grhl2* and *Tacstd2* expression, and inducing Trop2 expression in cysts (Fig. 1F, Fig. S1C).

These findings suggest that, in addition to causing a switch from multi-axial hepatocyte polarity to vectorial epithelial polarity (similar to bile duct cells), the silencing of Rab35, Dennd1b and Acap2 impaired hepatoblast-to-hepatocyte differentiation and increased the expression of genes characteristic of cholangiocyte identity.

### Isotropic expansion of apical lumina precedes transcriptional changes upon Rab35 silencing

The effects of silencing Rab35 or its partners on both the establishment of hepatocyte polarity and hepatoblast-to-hepatocyte differentiation, raised the question as to how these two processes are connected. Rab35 is an established regulator of cell polarity that plays a role in endocytic recycling (Klinkert and Echard, 2016). It is conceivable that Rab35 silencing affects the recycling of cell surface receptors involved in hepatoblast cell fate decisions. This could result in dysregulation of receptor-mediated signalling, driving the expression of cholangiocyte-related genes that lead to the formation of cyst-like lumina instead of BC. Alternatively, isotropic expansion of the apical lumen and/or subsequent changes in cell polarity phenotype could modulate a mechanotransduction pathway that feeds back on the cell transcriptional program (Meyer et al., 2020, Kaylan et al., 2018). To discern between these two scenarios, we examined the temporal dynamics of gene expression changes in relation to lumen formation throughout the 5-day culture of mock-, control (scrambled) siRNA-, and Rab35 siRNA-transfected cells (Fig. 2A, Table S2).

**Figure 2.**
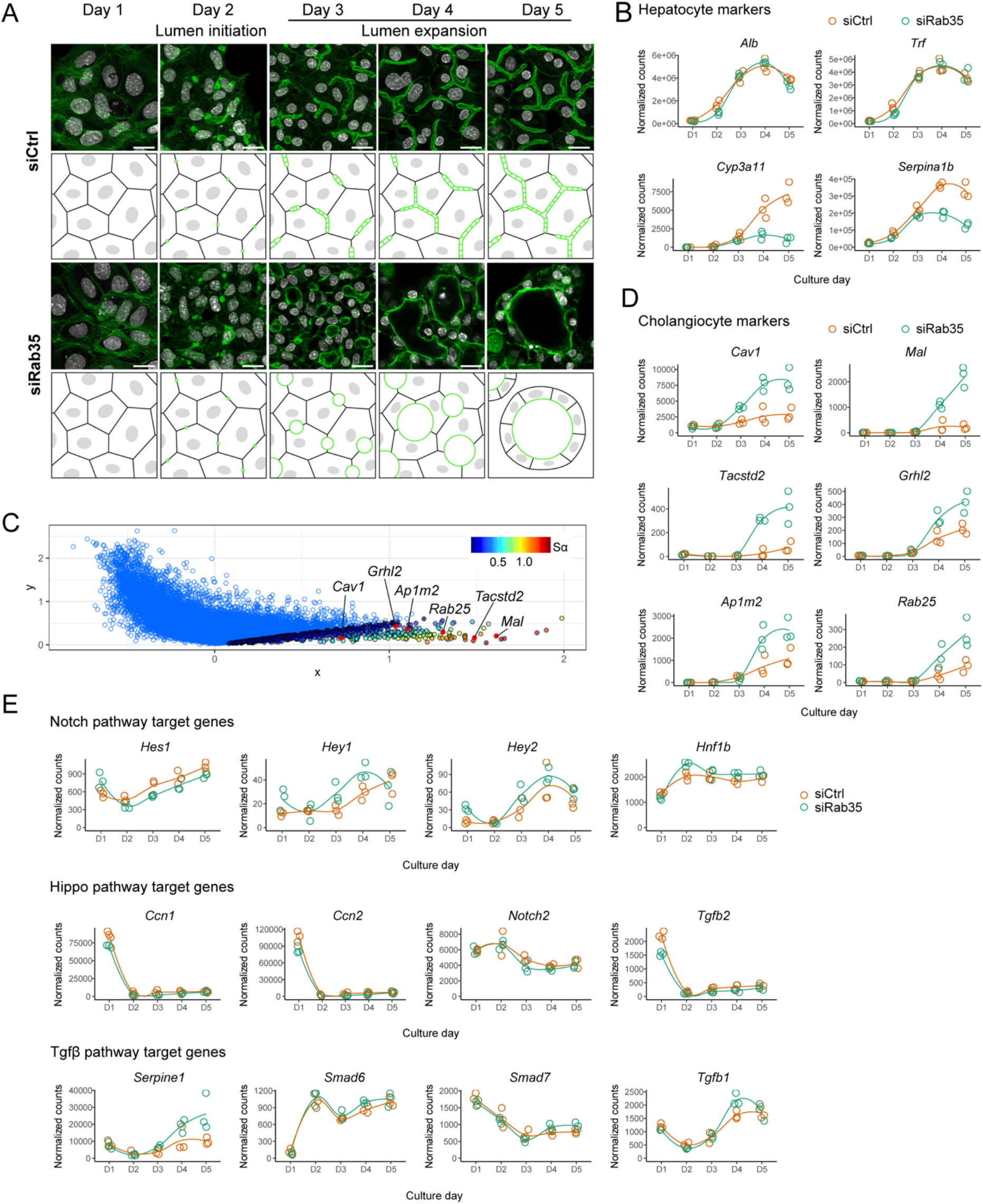
Isotropic lumen expansion precedes transcriptional changes following Rab35 silencing. A) Fluorescence microscopy and schematics showing *in vitro* hepatoblast-to-hepatocyte differentiation upon transfection with control and Rab35 siRNAs. F-actin (green), nuclei (grey). Scalebars: 20 μm. B) Expression profile of hepatocyte marker genes in siCtrl- and siRab35-transfected cells. Data points represent normalized counts of three biological repeats. C) Association Plot for the cluster of siRab35 day 4 and day 5 samples, generated using first five correspondence analysis dimensions. S_α_ of a gene is a ranking score of its cluster specificity. Cholangiocyte and vectorial polarity-related genes (e.g., *Mal, Tacstd2*) are highly specific for siRab35 day 4 and 5. D) Expression profile of genes associated with vectorial polarity and cholangiocyte identity in siCtrl- and siRab35-transfected cells. E) Expression profile of Notch, Hippo and Tgf-β targets in siCtrl- and siRab35-transfected cells.

Using the time-course dataset, we first confirmed hepatocyte differentiation in mock- and siCtrl-transfected cells. In both conditions, expression of the hepatoblast marker *Dlk1* decreased over time, while the expression of hepatocyte marker genes, such as *Alb, Trf, Cyp3a11*, and *Serpina1b*, increased, reflecting the differentiation of hepatoblasts into hepatocytes (Fig. S2A). Transfection with Rab35 siRNA resulted in silencing of Rab35 throughout the culture and *Dlk1* expression steadily decreased over time (Fig. S2B). Although Rab35 knockdown did not affect the expression of some hepatocyte markers, such as *Alb* and *Trf*, it clearly decreased the expression of others, e.g., *Cyp3a11* and *Serpina1b*, on day 4 and 5 (Fig. 2B, Fig. S2C)

To determine when gene expression between siCtrl- and siRab35-transfected cells starts to change, we visualized the complete time-course dataset using a correspondence analysis 3D biplot (File S1). Individual replicates of the samples clustered together, and the samples separated according to the culture timeline. Control and siRab35 samples were close to each other in the 3D plot on days 1-3, indicating that global gene expression patterns were similar during these days. The bifurcation between the siCtrl- and siRab35-transfected cells was observed at day 4, indicating notable differences in gene expression from day 4 onwards. Using an Association Plot (Fig. 2C), a two-dimensional plot that depicts genes associated to a cluster of samples (Gralinska and Vingron, 2023), we analysed genes characteristic for the Rab35 knockdown samples at days 4 and 5 and identified 588 genes in this cluster (cluster-specificity score cut-off: 0.01) (Table S3). Among those, we confirmed the upregulation of genes related to vectorial polarity and cholangiocyte identity, including *Ap1m2, Mal, Cav1, Rab25, Tacstd2* and the transcription factor *Grhl2* (Fig. 2D). Hence, the downregulation of hepatocyte-associated genes and upregulation of cholangiocyte-associated genes occurred simultaneously starting from day 4, i.e. one day after differences in lumen growth (anisotropic vs. isotropic expansion) emerge between siCtrl- and siRab35-transfected cells (Fig. 2A).

Previous work has demonstrated the role of Notch, Hippo and Tgf-β signalling in hepatoblast specification towards a cholangiocyte fate (Zong et al., 2009, Clotman et al., 2005, Yimlamai et al., 2014). To determine whether Rab35 knockdown stimulates the expression of cholangiocyte and vectorial polarity-associated genes by modulating these signalling pathways, we plotted the expression patterns of prototypical Notch, Hippo and Tgf-β target genes (Fig. 2E). None of the selected Notch and Hippo targets were altered by Rab35 knockdown and, except for Serpine1, Tgf-β targets were also unaffected. We thus conclude that modulation of canonical Notch, Hippo, or Tgf-β signalling is not the mechanism whereby Rab35 knockdown affects hepatocyte differentiation.

Interestingly, the time-course RNA sequencing experiment revealed that even mock- and siCtrl-transfected cells slightly increased the expression of various cholangiocyte-associated genes at days 4 and 5 (e.g., Fig. 2D, *Grhl2, Ap1m2, Tacstd2*). Indeed, at day 5 we occasionally observed Trop2-positive cells in control wells. Trop2 expression correlated with hepatocytes facing abnormally large apical lumina, typically found in regions with high cell density at the rims of the wells. (Fig. S3).

Together, the observation that the loss of apical bulkheads and consequent isotropic apical lumen expansion precede the transcriptional changes upon Rab35 silencing, and the expression of Trop2 in control cells with enlarged apical lumina, suggest a causal relationship between changes in apical lumen morphology (i.e., enlarged, more spherical, lumina instead of tubular BC) and impaired hepatocyte differentiation.

### Luminal pressure subapical actomyosin contractility influence hepatocyte differentiation

To test whether apical lumen morphology influences hepatocyte differentiation, we set to perturb lumen morphology using two methods, by reducing the contractility of the subapical actomyosin cortex and by increasing luminal pressure, and assessed the effect of these perturbations on the expression of cholangiocyte (*Tacstd2, Grhl2, Cav1, Ap1m2)* and hepatocyte (*Pck1* and *Cyp3a11*) marker genes.

First, to reduce the contractility of the subapical cortex, we silenced non-muscle myosin IIa (NMIIA, *Myh9*), which is the most abundant non-muscle myosin isoform in differentiating hepatoblasts and localizes to the subapical actomyosin cortex (Fig. 3A,B). Similar to Rab35 silencing, knockdown of NMIIA caused isotropic apical lumen expansion, and ultimately the reorganization of hepatocytes into cysts with vectorial polarity (Fig. 3C,D). Furthermore, NMIIA knockdown reduced *Cyp3a11* and *Pck1* expression, while it increased *Ap1m2* and *Grhl2* expression (Fig. 3E).

**Figure 3.**
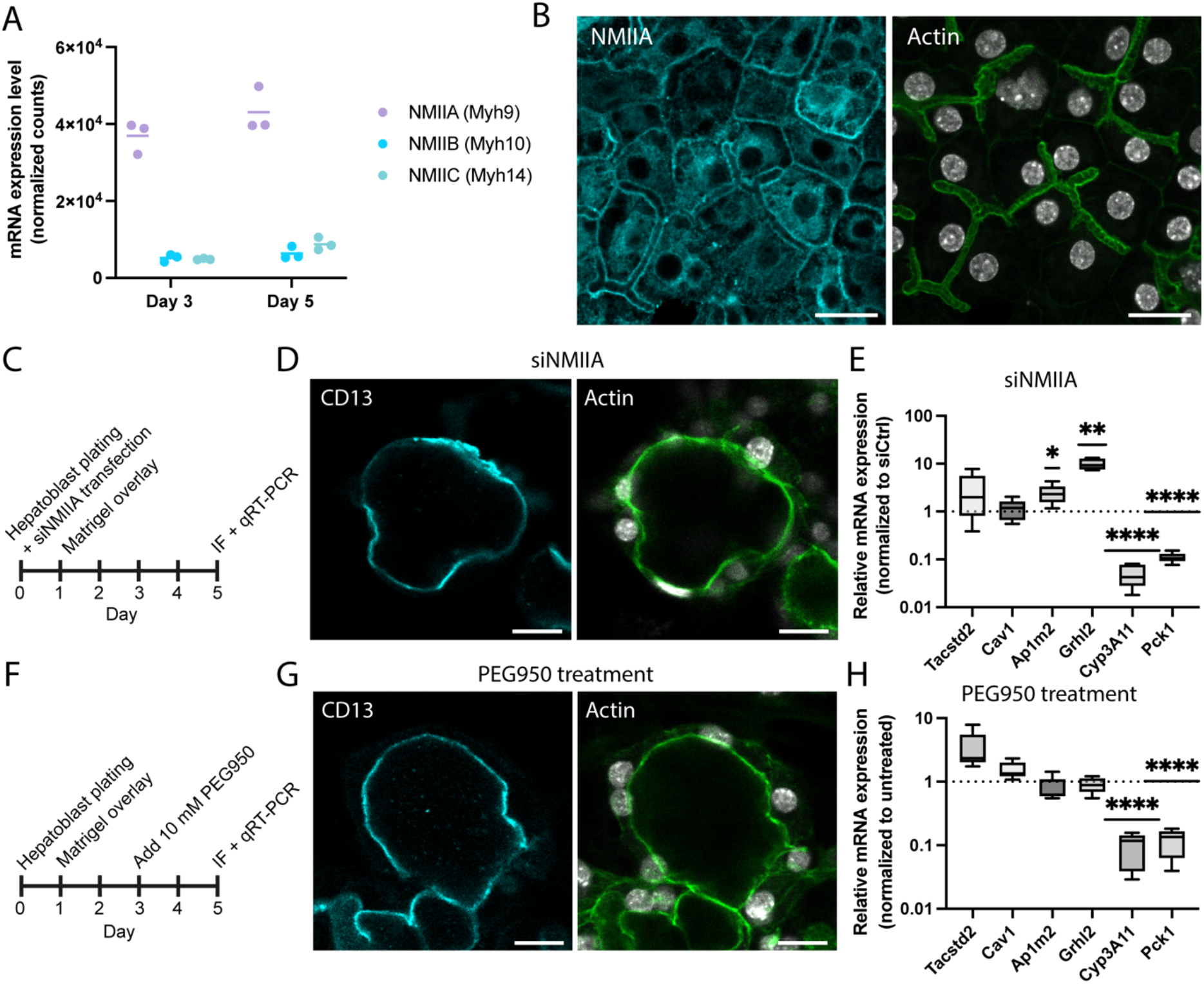
Modulation of actomyosin contractility and luminal pressure impairs hepatocyte differentiation. A) Expression levels of non-muscle myosin (NMII) isoforms in siCtrl-transfected hepatoblasts at day 3 and 5 based on RNA sequencing. B) Differentiating hepatoblasts at day 5. NMIIA (cyan), F-actin (green) and nuclei (grey). C) Timeline of the NMIIA knockdown experiment. IF: Immunofluorescence microscopy D) siNMIIA-transfected differentiating hepatoblasts. CD13 (cyan), F-actin (green), nuclei (grey). E) qRT-PCR for genes associated with vectorial polarity and cholangiocyte identity (*Tacstd2, Cav1, Ap1m2, Grhl2*) and hepatocyte identity (*Cyp3a11, Pck1*) after NMIIA knockdown. F) Timeline of the PEG950 treatment. G) PEG950-treated differentiating hepatoblasts. CD13 (cyan), F-actin (green), nuclei (grey). H) qRT-PCR for genes associated with vectorial polarity and cholangiocyte identity (*Tacstd2, Cav1, Ap1m2, Grhl2*) and hepatocyte identity (*Cyp3a11, Pck1*) upon PEG950 treatment. Box plots represent 5 independent experiments. *, P < 0.05; **, P < 0.01; ****, P < 0.0001. Scalebars: 20 μm.

Second, we raised luminal pressure by treating the cells with 10 mM polyethylene glycol 950 (PEG950). PEG950 is an osmotically active biologically inert polymer that is transported into the BC lumen (Javitt, 2014), resulting in water influx into the BC and elevated hydrostatic pressure. PEG950 treatment between day 3 and 5 induced the formation of cysts with vectorial polarity (Fig. 3F,G). Moreover, this approach led to reduced levels of *Cyp3a11* and *Pck1* and slightly elevated levels of *Cav1* and *Tacstd2* (Fig. 3H).

The similarity of the response to reduced actomyosin contractility and increased hydrostatic pressure suggests the existence of a mechanotransduction pathway that leads to transcriptional changes. However, whereas both NMIIA knockdown and PEG950 treatment reduced the expression of the two hepatocyte markers, they only partially recapitulated the upregulation of the cholangiocyte-associated genes as observed upon Rab35 knockdown (significant upregulation of *Ap1m2* and *Grhl2* upon NMIIA knockdown, increased expression of *Cav1* and *Tacstd2* upon PEG950 treatment). Rab35 participates in multiple cellular processes, such as actin cytoskeleton organization and endocytic recycling (Klinkert and Echard, 2016). The upregulation of cholangiocyte-associated genes upon Rab35 knockdown likely results from the loss of multiple Rab35-dependent processes that affect both luminal pressure and actomyosin contractility.

In conclusion, our findings suggest that the induction of isotropic apical lumen expansion, instead of tubular BC elongation, impairs hepatoblast-to-hepatocyte differentiation and maturation, demonstrated by the decrease in hepatocyte markers and the upregulation of genes associated with cholangiocyte fate. The specific mechanotransduction pathways that mediate these transcriptional changes upon loss of lumen anisotropy are currently unknown. The Notch and Hippo signalling pathways are sensitive to mechanical cues and implicated in cholangiocyte specification (Dupont et al., 2011, Gordon et al., 2015, Hunter et al., 2019, Zong et al., 2009, Yimlamai et al., 2014). However, the absence of transcriptional changes in the main Notch and Hippo target genes (Fig. 2E) suggests that canonical Notch and Hippo signalling pathways do not cause the changes in gene expression upon Rab35 knockdown.

### The interplay of cell polarity and cell fate decisions in the developing liver

During liver development, most hepatoblasts differentiate into hepatocytes through a default program mediated by progressively strengthened gene regulatory networks of hepatocyte transcription factors (Yang et al., 2017, Kyrmizi et al., 2006, Yang et al., 2023). A small proportion of hepatoblasts receives signals from the portal mesenchyme to take on a cholangiocyte fate and form bile ducts. This process involves several signalling pathways, e.g. Notch, Hippo and Tgf-β, which drive the expression of cholangiocyte-associated transcription factors, e.g. Sox9, Hnf1β (Zong et al., 2009, Clotman et al., 2005, Yimlamai et al., 2014, Lee et al., 2016, Wu et al., 2017). In this study, we found an additional level of cell fate regulation that integrates control of gene expression with cell polarity and lumen morphogenesis, adding to the body of evidence that the establishment of epithelial polarity is not merely an endpoint to cell differentiation, but actively contributes to cell fate decisions (Motegi et al., 2020). We showed that the failure to establish the typical elongated BC results in the generation of cysts, where the cells assume a cholangiocyte-like vectorial polarity, have reduced expression of hepatocyte-specific genes and increased expression of cholangiocyte-associated genes (Fig. 4). We termed these cells vectorial hepatocytes, as they do not seem to acquire a complete cholangiocyte fate upon Rab35 silencing *in vitro* (studied here), or in embryonic livers *in vivo* (Belicova et al., 2021), as evidenced by the expression of Hnf4α and lack of Sox9 expression. Nevertheless, the expression of cholangiocyte-specific genes indicates that these cells assume features of both cell types. It would be interesting to explore if these cells eventually fully differentiate into cholangiocytes, i.e. express a full set of cholangiocyte markers, establish cilia, suppress Hnf4α, etc. Unfortunately, the hepatoblasts in our primary culture are only viable for up to 6 days, which may be too short to follow whether they could complete cholangiocyte differentiation.

**Figure 4.**
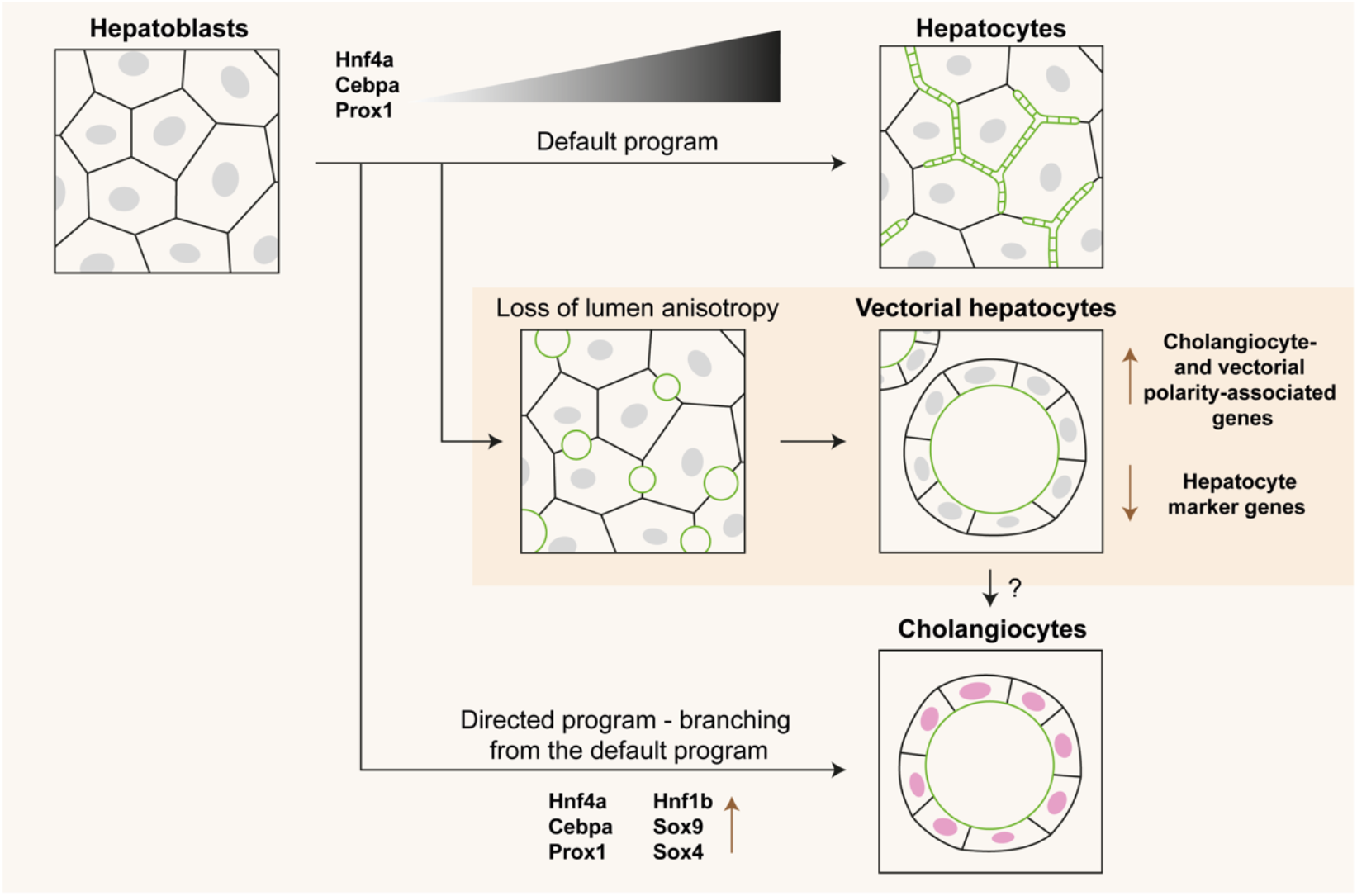
The consequence of a loss in lumen anisotropy on hepatoblast differentiation.

*In vivo*, bile ducts develop through a stage of transient asymmetry where one side is composed of Sox9-positive cholangiocytes and the other side of HNF4α-positive hepatoblasts (Antoniou et al., 2009). These hepatoblasts acquire vectorial polarity, contribute to an expanding multicellular lumen, and eventually differentiate into cholangiocytes as the bile duct matures. Based on our results, we propose that the acquisition of vectorial polarity is an additional regulatory step in the differentiation of hepatoblasts towards cholangiocytes in asymmetric bile ducts. Additionally, apical lumen morphology and hepatocyte polarization may influence cell fate in the context of liver disease. Cholestatic liver diseases are often associated with the generation of so-called hepatocyte rosettes. In these structures, hepatocytes are vectorially polarized towards a common enlarged apical lumen, and some cells do express cholangiocyte markers, such as Sox9 (Lin et al., 2022, Mayer et al., 2023). This suggests that the feedback of cell polarity and lumen formation on cell (de)differentiation described here may operate under pathological conditions as during hepatocyte rosette formation.

In conclusion, our data demonstrates that anisotropic BC elongation is required for hepatoblast-to-hepatocyte differentiation. Further studies into the mechanisms that link cell polarity, morphogenesis, and cell fate decisions in the liver are highly relevant to understand how liver tissue can be resilient to changes in luminal pressure during development and disease.

## Materials and methods

### Hepatoblast isolation and in vitro differentiation toward hepatocytes

Hepatoblast isolation from embryonic livers (E13.5) of C57BL/6JOlaHsd mice and subsequent *in vitro* differentiation toward hepatocytes was performed as described in Belicova et al. (2021). In brief, embryonic livers were fragmented in Liver Perfusion Medium (Thermo Fisher Scientific; 17701–038) and subsequently digested in Liver Digest Medium (Thermo Fisher Scientific; 17703–034) supplemented with 10 µg/ml DNase I (Sigma-Aldrich; DN25). Erythrocytes were lysed in red blood cell lysis buffer: 155 mM NH4Cl, 10 mM KHCO3, and 0.1 mM EDTA, pH 7.4. Next, the cells were blocked with anti-mouse CD16/CD32 blocking antibody (BD Biosciences; 553142; Rat) and Dlk1+ hepatoblasts were labeled with anti-Dlk mAb-FITC (MBL; D187-4; Rat) and anti-FITC MicroBeads (Miltenyi Biotec; 130–048-701). Labeled Dlk + hepatoblasts were isolated using a magnetic column (Miltenyi Biotec; 130–024-201) according to the manufacturer’s protocol. Isolated hepatoblasts were seeded in 96-well cell culture microplates (Greiner; 655090) coated with Matrigel (Corning; 356231) at a density of 13,000 cells/well in expansion media: DMEM/F-12, GlutaMAX supplement (Thermo Fisher Scientific; 31331028), 10% FBS, 1 × insulin-transferrin-selenium-ethanolamine (Gibco; 51500–056), 0.1 µM dexamethasone (Sigma-Aldrich; D1756), 10 mM nicotinamide (Sigma-Aldrich; N0636), 10 ng/ml human hepatocyte growth factor (in-house production), and 10 ng/ml mouse epidermal growth factor (in-house production). After 24 h, the cells were overlaid with differentiation media: MCDB131 (Thermo Fisher Scientific; 10372019), 2 mM L-glutamine (Thermo Fisher Scientific; M11-004) 0.25 × insulin-transferrin-selenium-ethanolamine, and 0.1 µM dexamethasone, containing Matrigel to a final concentration of 5% vol/vol. Cells were cultured in a humidified incubator with 5% CO2 at 37°C for a maximum of 6 days. For the PEG950 treatment, the culture medium of the cells was replaced at day 3 with differentiation medium containing 10 mM PEG950 (Merck; P3515).

### siRNA-mediated knockdown

Cells were reverse transfected with siRNAs upon plating using Lipofectamine RNAiMAX (Thermo Fisher Scientific; 13778075) according to the manufacturer’s protocol. The final siRNA concentration per well was 10 nM (siRab35, siNMIIA) or 20 nM (siAcap2, siDennd1b) at a concentration of 0.1 vol/vol % Lipofectamine RNAiMAX. siluciferase was used as control for siAcap2, siDennd1b and siNMIIA. A scrambled Rab35 siRNA (scrRab35) was used as control for siRab35 in all experiments, except for the RNA sequencing experiment in Fig. 1, where siLuciferase was used as control. siRNAs used: siLuciferace (sense 5′-cuuAcGcuGAGuAcuucGAdTsdT-3′, antisense 5′-UCGAAGuACUcAGCGuAAGdTsdT-3′), siNMIIA (Myh9) (Thermo Fisher Scientific; MSS237342), siRab35 (sense: 5’-CGcAAGGAGGAGcAUUUuATsT-3’, antisense: 5’-uAAAAUGCUCCUCCUUGCGTsT-3’), scrRab35 (sense: 5’-GuAcGuAAucGGcAGAAGuTsT-3’, antisense: 5’-ACUUCUGCCGAUuACGuACTsT-3’), siDennd1b (sense: 5’-AuGAGAGGcGcAucAuuAuTsT–3’, antisense: 5’-AuAAUGAUGCGCCUCUcAUTsT-3’) and siAcap2 (sense: 5’-cAAuGuGcuucAGucAAAATsT-3’, antisense: 5’-UUUUGACUGAAGcAcAUUGTsT-3’).

### Immunofluorescence staining and confocal microscopy

Cells were fixed with 3% PFA in PBS for 20 min at RT, washed with PBS, and blocked using blocking buffer (2% BSA and 0.05% saponin in PBS) for 1h at RT. Cells were incubated with primary antibodies diluted in blocking buffer at 4°C overnight. Primary antibodies used: anti-CD13 (NB100-64843; Rat; Novus), anti-NMIIA (GTX113236; Rabbit; Genetex), anti-Trop2 (ab214488; Rabbit; Abcam) and anti-Caveolin (D46G3; Rabbit; Cell Signaling). Phalloidin-488 (A12379; Thermo Fisher Scientific) and DAPI were included in the primary antibody solutions. Following washing with PBS, cells were incubated with secondary antibodies diluted in blocking buffer for 2h at RT. Secondary antibodies used were goat anti-rabbit-AlexaFluor-647 (A21244; Thermo Fisher Scientific) and goat anti-rat-AlexaFluor-647 (A21247; Thermo Fisher Scientific). An LSM700 inverted laser scanning confocal microscope (Zeiss) equipped with a 40x/1.2 C-Apochromat, water, DIC objective (Zeiss), and Zen imaging software (Zeiss) were used for confocal microscopy. Fiji (Schindelin et al., 2012) was used for image analysis.

### RNA isolation and RT–quantitative PCR (qPCR)

RNA was isolated using the RNeasy Mini Kit (Qiagen; 74104) including the DNase I (Qiagen; 79254) treatment. Cells were lysed with RNeasy lysis buffer supplemented with DTT. cDNA was synthesized using the ProtoScriptII First Strand cDNA Synthesis Kit (NEB; E6560S), following the manufacturer’s protocol with the Random Primer Mix and the RNA denaturation step. qPCR was performed with the Roche LightCycler 96 using FastStart Essential DNA Green Master (Roche; 06402712001). As reference gene we used the housekeeping gene Rplp0. Primers sequences used are described in Table 1. Normalized relative gene expression value and percent knock-down was calculated using the ΔΔCq method (Haimes and Kelley, 2010). GraphPad Prism version 9.0 (GraphPad software) was used for data analysis and visualization.

**Table 1.**
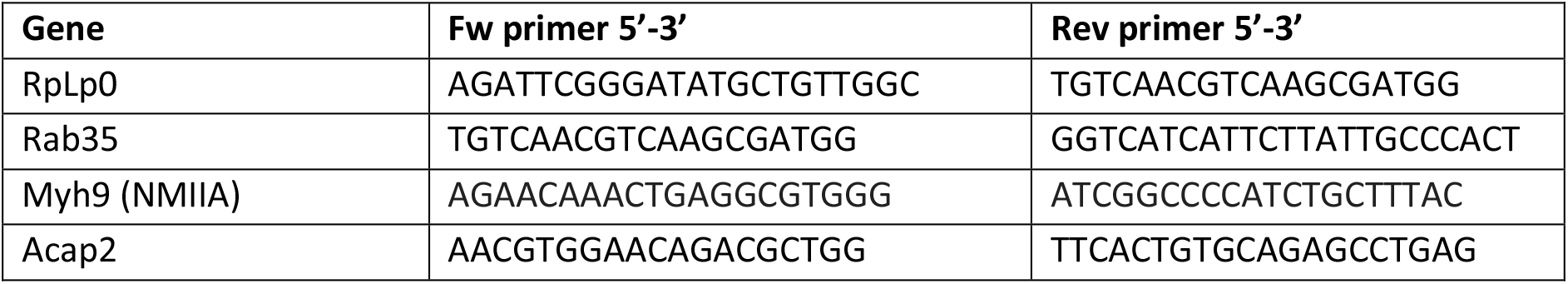
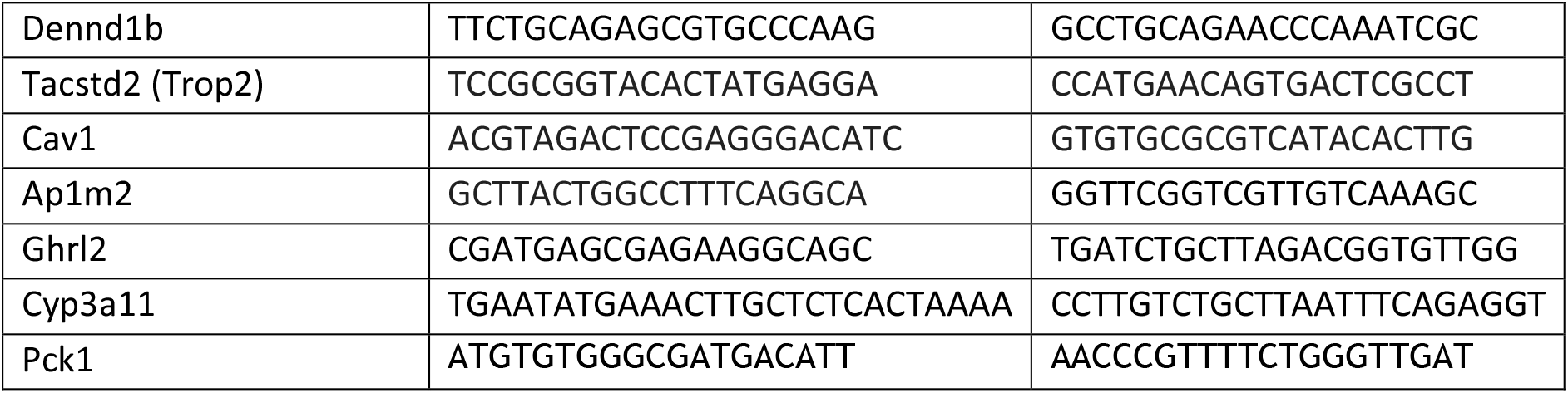

### RNA sequencing

For the RNA sequencing experiment in Figure 1B, samples were collected from 3 biological replicates of in vitro cultured E13.5 Dlk+ hepatoblasts transfected with siRNA targeting Rab35 (siRNA #4), or non-targeting control Luciferase (siLuc) at day 5 of the culture. For the time-course RNA sequencing of Figure 2, samples were collected every day from day 1 to day 5 from Mock-, scrRab35-, or siRab35-transfected in vitro cultured E13.5 Dlk+ hepatoblasts in 3 biological replicates. Rab35 mRNA knockdown was verified by RT-qPCR. The integrity of RNA was measured by Agilent 2100 Bioanalyzer. Preferentially, only samples with the RNA integrity number (RIN) > 9.0 were used. mRNA was isolated from the total RNA by poly-dT enrichment using the NEBNext Poly(A) mRNA Magnetic Isolation Module according to the manufacturer’s instructions, fragmented to 350 nt, and libraries were prepared using the NEBNext UltraTM II Directional RNA Library Prep Kit for Illumina. For Illumina flowcell production, samples were equimolarly pooled and distributed on all lanes used for 75bp single read sequencing on Illumina NextSeq 500 (Fig. 1B) or 100bp paired-end sequencing on Illumina NovaSeq 6000 (Fig. 2), resulting in an average of 33 Mio and 67 Mio sequenced fragments per sample, respectively.

Analysis of RNA sequencing data was performed in R (v4.1.0) (R Core Team, 2021) using following packages: DESeq2 (v1.34.0) (Love et al., 2014), ggplot2 (v3.4.0) (Wickham, 2009), pheatmap (1.0.12) (Kolde, 2019), EnhancedVolcano (v1.12.0) (Blighe et al., 2021), VennDiagram (v1.7.0) (Chen, 2021), dplyr (1.0.7) (Wickham et al., 2021), ggpubr (v0.5.0) (Kassambara, 2022), cowplot (v1.1.1) (Wilke, 2020). Online application ShinyGo (v0.741) (Ge et al., 2020) was used for Gene ontology enrichment analysis (KEGG pathways database). DESeq2 package was used with the default parameters (e.g., Benjamini-Hochberg correction for p-value). The replicates in both experiments clustered well together and batch correction was not required. Correspondence analysis (CA) of the time-course RNA sequencing data was conducted using the Bioconductor package APL (v0.99.5) (Gralinska et al., 2022). The function cacomp was applied to the matrix of normalized counts (output of the DESeq2 package) using the default parameters, except for the top set to 19331. The interactive 3D plot was generated using Plotly (Plotly Technologies Inc.) (Sievert, 2020). Association Plot for the cluster of Rab35 siRNA-treated samples from days 4 and 5 was generated using the first five CA dimensions. Cluster-specificity scores Sα were computed based on 500 permutations (parameter reps in function apl_score).

In the time-course RNA sequencing experiment, scrRab35-transfected cells showed a delay in both lumen formation and differentiation compared to mock-transfected cells, e.g., day 4 siRNA-treated samples clustered with day 3 mock-transfected cells (Fig. S4A). Therefore, we used the scrRab35 condition as the appropriate control to siRab35-transfected cells. Importantly, even though different control siRNAs were used (siLuc vs. scrRab35), the results of the RNA sequencing experiment on day 5 (Fig. 1B) were comparable to the results of the time-course RNA sequencing experiment on day 5 (Fig. S4B,C), indicating that the results were reproducible.

### Proteomics

On day 6 of the hepatoblast culture, cells were washed twice with PBS at RT, placed on ice and washed 3 times 10min with cell recovery solution (Corning; CLS354270) to remove Matrigel. Cells were lysed at RT for 10 min using freshly prepared lysis buffer: RIPA buffer with EDTA and EGTA (Alfa Aesar/Thermo Fisher Scientific) + 5% SDS + 2x cOmplete protease inhibitor cocktail (Roche). After centrifugation at 14k rpm for 10 min at RT, supernatant was used for proteomics. Label-free protein quantification was performed with 4 biological replicates per condition, prepared and measured in a block-randomised fashion (Burger et al., 2021). After bead beating with a TissueLyser II (QIAGEN) and 0.5 mm stainless steel beads (Next Advance) for 2 rounds of 5 min at 25 Hz at 4 degrees to sheer DNA/RNA, samples were centrifuged at 13k g for 10 min at RT. Proteins from the supernatants were precipitated in final 85% acetone and 30 mM NaCl by incubation at RT for 30 min at 800 RPM in a shaker (Nickerson and Doucette, 2020). Protein pellets were resuspended in 1x S-Trap lysis buffer (5% SDS, 50 mM TEAB from Merck, pH 8.5). 50 µg of the sample protein was digested using the S-trap micro spin column digestion protocol from ProtiFi LLC. Washes were performed 6x with 200 µL each and digestion was performed with 0.75 µg Trypsin/Lys-C Mix (Promega) per 10 µg total protein. Eluted peptides were dried in a RVC 2-25 rotary vacuum concentrator (Martin Christ Gefriertrocknungsanlagen GmbH), and stored at -20 °C. Before measurement, samples were resuspended in 0.2% formic acid (Merck). Total peptide concentration was adjusted to 0.15 µg/µL according to the absorbance at 280 nm against a Mass Spec-Compatible Human Protein Extract/Digest calibration curve (Promega), using the Thermo Nanodrop 1000 ND-1000 spectrophotometer (Thermo Fisher).

The LC was a Thermo Dionex UltiMate 3000 UHPLC system with an Acclaim PepMap precolumn (100 µm x 20 mm, C18, 5 µm, 100 Å) and a 50 cm µPAC column array (all from Thermo Fisher). 5 µl of the sample was loaded onto the precolumn at 5 µl/min and eluted with a linear gradient with 2 slopes at 0.5 µl/min and 0.1 % formic acid. The percentage of acetonitrile (ACN) (Thermo Fisher) was increased from 0 to 17.5 % over two-thirds of the gradient length and from 17.5 % to 35 % in the last third. Both columns were then rinsed with 95 % ACN for 7 minutes. At least one blank injection with a 25-minute gradient was performed between each sample injection to reduce carryover. Sample measurements with 120-minute gradients were performed. We used a μPAC Flex iON interface (Pharma Fluidics / Thermo Fisher) equipped with a 5 cm nanoESI emitter with a 20 µm inner diameter (FOSSILIONTECH). Electrospray was performed with a spray voltage of 2.5 kV, a capillary temperature of 280 degrees Celsius, and an S-lens RF value of 50.

MS data acquisition was performed using a Q Exactive HF mass spectrometer (Thermo Fisher Scientific, Bremen, Germany). Data-independent acquisition included a low-resolution MS1 scan in centroid mode covering the *m/z* range of 395-971 with a resolution R*m/z* =200 of 30,000, an AGC of 3 × 10^6^, 40 ms injection time, and a fixed first mass at *m/z* 100. The following 32 MS2 scans covered the *m/z* range of 400 to 966 (after demultiplexing) with a total cycle time of about 3 seconds. MS2 spectra were acquired under normalized collision energy (NCE) of 24 using an *m/z* isolation window width of 18, maximum filling time of 55 ms under AGC of 1× 10^6^, and resolution R_m/z=200_ of 30,000 in centroid mode. DIA acquisition was performed in a staggered manner (Amodei et al., 2019). The raw files were converted to mzML format and demultiplexed with MSConvert v3.0.2 (Adusumilli and Mallick, 2017). Analysis against mouse reference proteome UP000000589 SwissProt canonical+isoform as of 07.02.2022 with added MaxQuants common contaminants was performed with DIA-NN v1.8 (Demichev et al., 2020). Settings were --min-fr-mz 100 --max-fr-mz 2000 --cut K*,R* --missed-cleavages 1 --min-pep-len 7 --max-pep-len 34 --min-pr-mz 400 --max-pr-mz 966 --min-pr-charge 2 -- max-pr-charge 4 --var-mods 1 --var-mod UniMod:35,15.994915,M --double-search --individual-mass-acc --individual-windows --smart-profiling --pg-level 2 --species-genes --peak-center --no-ifs-removal - -no-quant-files --report-lib-info --matrix-qvalue 0.01 --nn-single-seq --fixed-mod UniMod:39,45.987721,C --strip-unknown-mods. For further analysis, the protein group matrix from DIA-NN was processed. Protein groups reported in at least 2 out of the total of 4 replicates per condition were considered as positively identified. Protein groups further characterized by a CV (coefficient of variation) below 25% were considered as quantified in that condition. Differential expression analysis of quantified protein groups was performed with Limma 3.50.0 with α = 0.01 and ∣log2 fold-change∣ > 0.5 (Ritchie et al., 2015). DIA data acquisition and DIA-NN data processing routine were optimized and validated according to (Jumel and Shevchenko, 2024)

## Statistical analysis

qRT-PCR experiments in Fig 1F and Fig 3E,H were analyzed using Student’s one-sample two-tailed t-test against a theoretical mean of 1.

## Data availability

Raw data will be provided upon reasonable request. Raw data will be deposited into data repositories once the final article is published.

## Acknowledgements

We thank Meritxell Huch for valuable feedback and critical reading of the manuscript. We acknowledge the light microscopy facility, the biomedical services facility, and André Gohr and Lena Hersemann from the scientific computing facility at Max Planck Institute of Molecular Cell Biology and Genetics in Dresden for their contributions. We acknowledge the DRESDEN-concept Genome Center for the bulk RNA-seq service.

This work was financially supported by the European Molecular Biology Organization (grant. no. ALTF 509-2021) to M.P. Bebelman, the German Federal Ministry of Research and Education (LiSyM-Krebs, grant no. 031L0258C) and the European Research Council ERC Advanced Grant (no. 695646) to M. Zerial. Open Access funding provided by the Max Planck Society.

## Author contributions

Conceptualization: L. Belicova, M. Bebelman, and M. Zerial. Methodology: L. Belicova, M. Bebelman, T. Jumel, A. Lahree. Investigation: L. Belicova, M. Bebelman, T. Jumel.

Software: E. Gralinska, L. Belicova, Tobias Jumel.

Formal analysis: L. Belicova, M. Bebelman, E. Gralinska, Tobias Jumel. Resources: T. Zatsepin, M. Vingron, Andrej Shevchenko.

Visualization: L. Belicova, E. Gralinska, M. Bebelman. Validation: L. Belicova; M. Bebelman.

Writing – original draft: L. Belicova, M. Bebelman.

Writing – review and editing: L. Belicova, M. Bebelman, E. Gralinska, A. Lahree, T. Jumel, Y. Kalaidzidis, M. Zerial.

Supervision: M. Zerial, Y. Kalaidzidis, M. Vingron, Andrej Shevchenko. Funding acquisition: M. Bebelman, M. Zerial.

---

**Figure S1.**
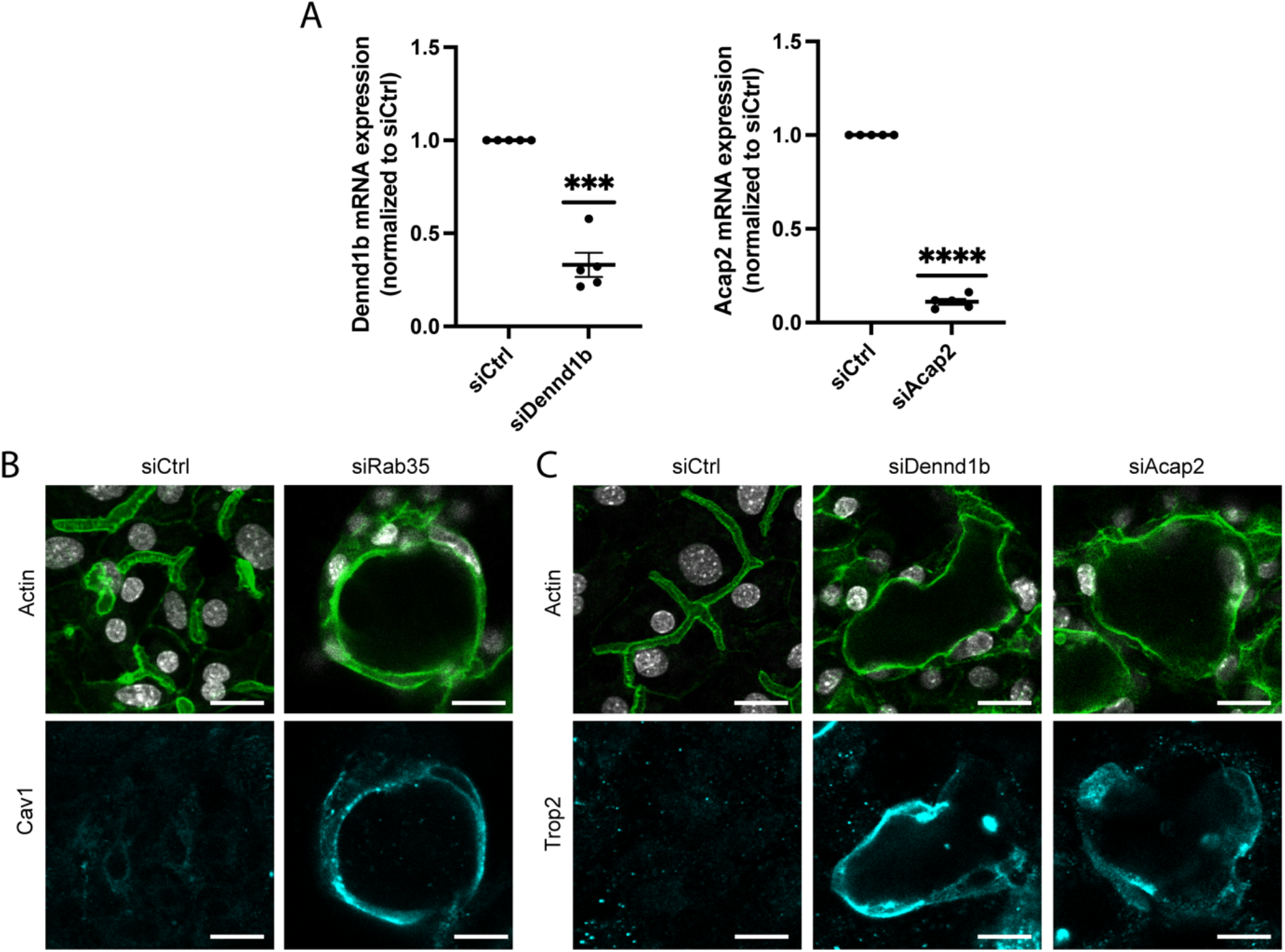
A) qRT-PCR analysis showing knockdown of Dennd1b and Acap2 at day 5 of culture following transfection with Dennd1b and Acap2 siRNA, respectively. Graph represents data from 5 independent experiments. ***, P < 0.001; ****, P < 0.0001 using Student’s one-sample two-tailed t-test. B) Immunofluorescence microscopy of differentiating hepatoblasts transfected with control or Rab35 siRNA at day 5 of culture. Cav1 (cyan), F-actin (green) and nuclei (grey). Scalebar: 20 μm. C) Immunofluorescence microscopy of differentiating hepatoblasts transfected with control, Dennd1b or Acap2 siRNA at day 5 of culture. Cells are stained for Trop2 (cyan), F-actin (green) and nuclei (grey). Scalebar: 20 μm.

**Figure S2.**
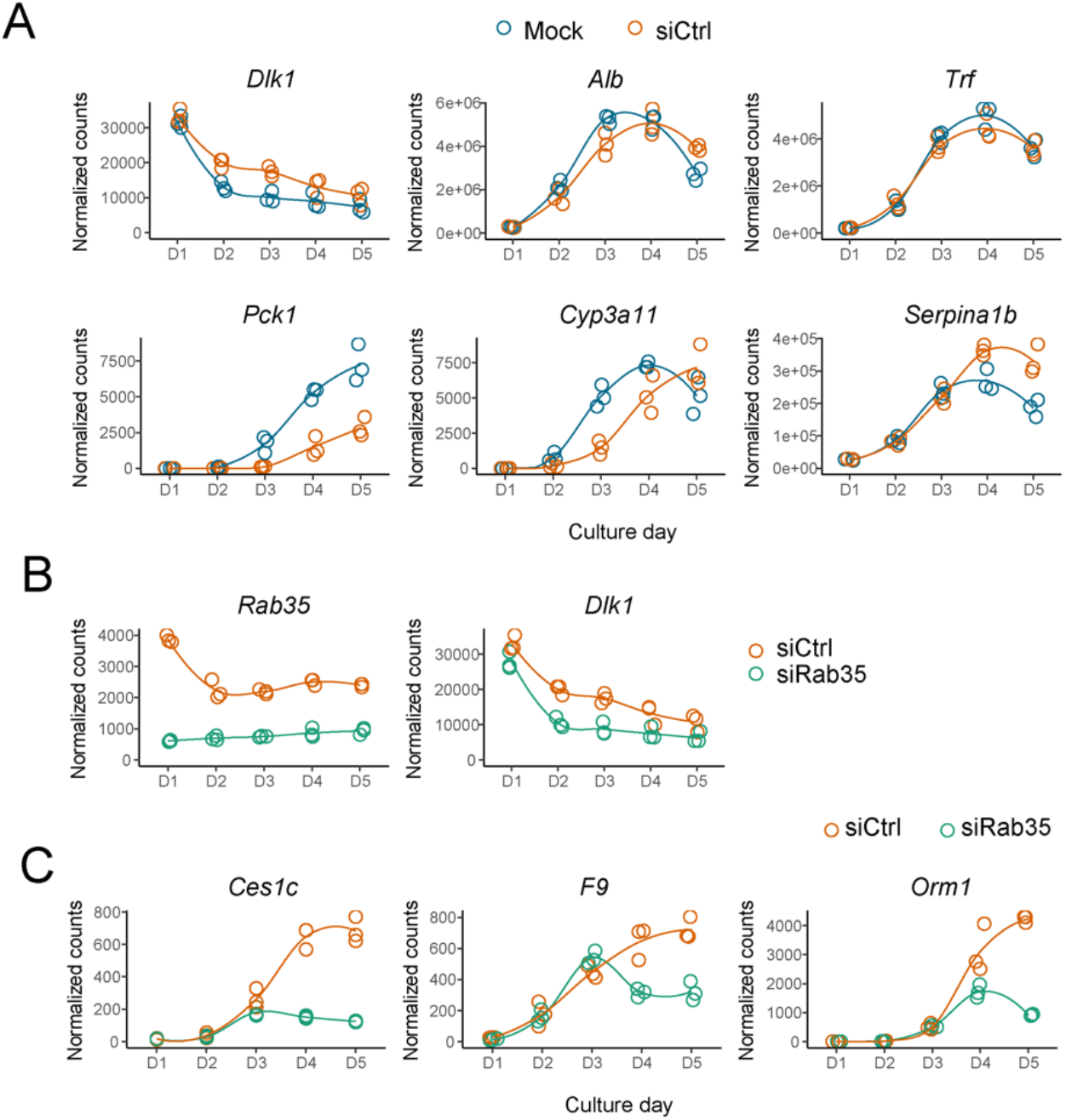
A) Expression profile of hepatoblast or hepatocyte marker genes in mock and control (scrambled) siRNA transfected cells at each day of the culture based on RNA sequencing. Data points represent normalized counts of three biological repeats. siCtrl data was also used to compare to siRab35 in Figure 2B and Figure S2B. B) Expression profile of *Rab35* and *Dlk1* (hepatoblast marker) in control and Rab35 siRNA (siRab35) treated samples, three biological replicates represented by data points. C) Expression profile of hepatocyte markers in control and Rab35 siRNA (siRab35) treated samples, three biological replicates represented by data points.

**Figure S3.**
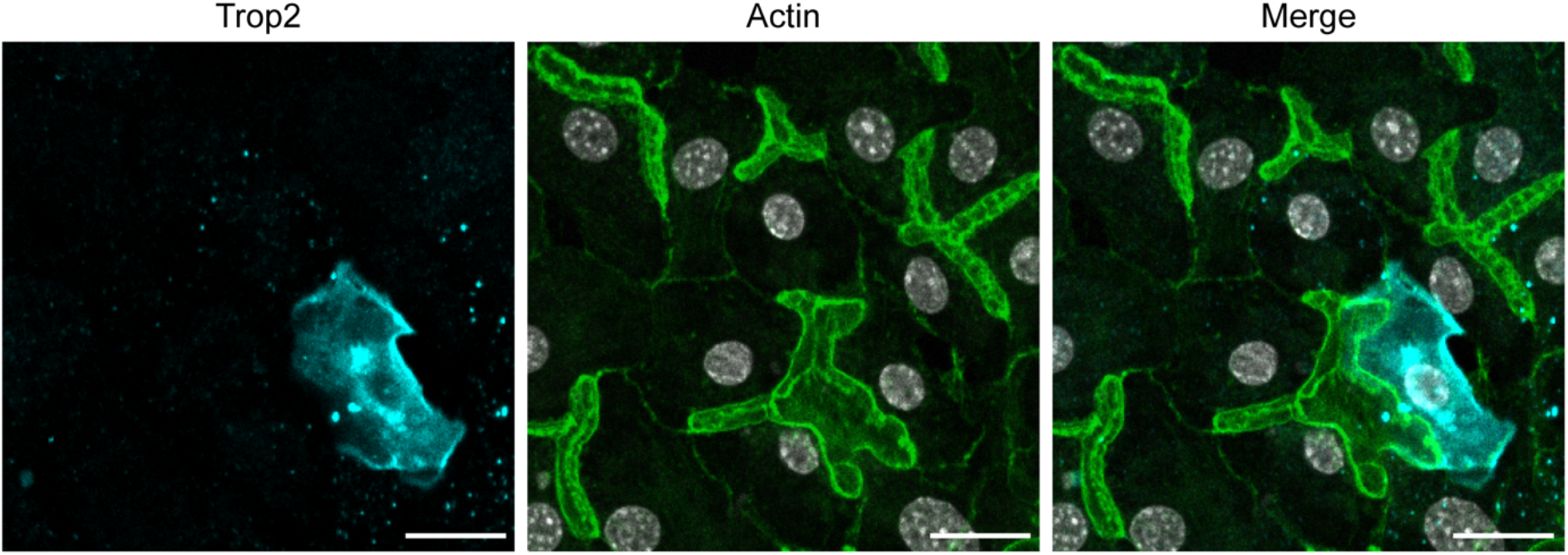
Immunofluorescence microscopy of differentiating hepatoblasts transfected with control siRNA (siLuciferase) at day 5 of culture. Cells are stained for Trop2 (cyan), F-actin (green) and nuclei (grey). Scalebar: 20 μm.

**Figure S4.**
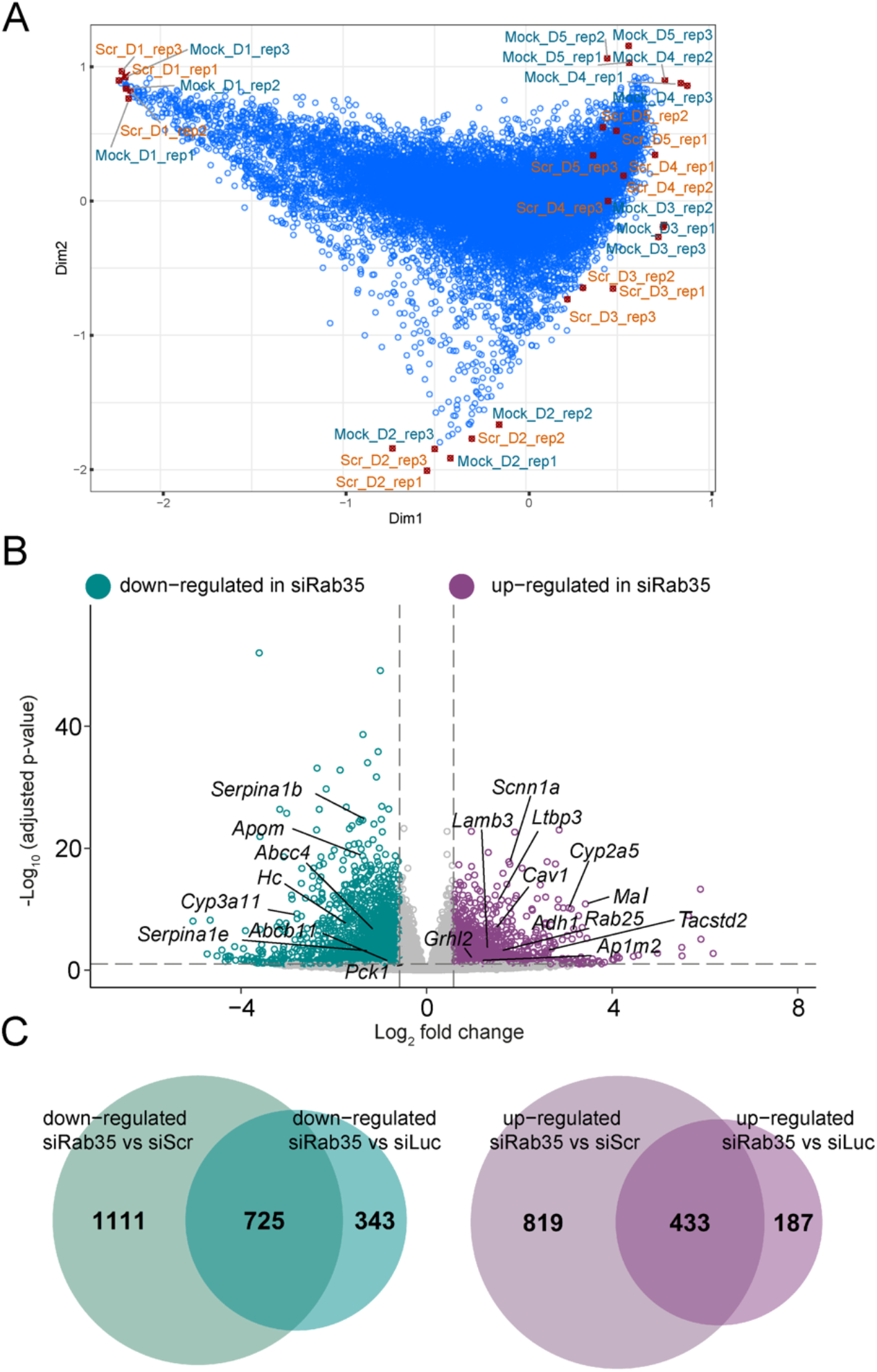
A) Correspondence analysis biplot of time-course (D1 -D5) mock and control (Scr) siRNA bulk RNA-seq data. Blue dots represent expressed genes, while red dots individual replicates of the samples. B) Volcano plot of results from differential gene expression analysis between control (scrambled) and Rab35 siRNA (siRab35) treated cells at day 5 of the time-course experiment. Significant genes (log_2_ fold change > |0.58|, p-adjusted value < 0.1) in green are down-regulated in siRab35 samples and in magenta up-regulated. The labelled genes are the same as in Fig. 1B, where siRab35 samples were analysed in comparison to Luciferase siRNA control samples. C) Venn diagram visualizing the number of genes identified as down-regulated or up-regulated (log_2_ fold change > |0.58|, p-adjusted value < 0.1) in siRab35 samples compared to siRNA control at day 5 in experiment from Fig. 1B and from the time-course experiment in panel E, and their overlap.

